# SALSA: resolving sign ambiguity in M/EEG source reconstruction with no task dependence

**DOI:** 10.64898/2026.11.16.766397

**Authors:** Ryshum Ali, Olivier Burta, Chiara Rossi, Mats W.J. van Es, Chetan Gohil, Mark W. Woolrich, Jeroen Van Schependom, Diego Vidaurre Henche

**Affiliations:** Epilepsy Center, Department of Neurology, University Hospital Erlangen, Friedrich-Alexander-University Erlangen-Nürnberg (FAU), Erlangen, Germany; AIMS lab, Vrije Universiteit Brussel, Center for Neurosciences, Brussels, Belgium; Department of Electronics and Informatics (ETRO), Vrije Universiteit Brussel, Brussels, Belgium; Oxford Centre for Human Brain Activity, Oxford Centre for Integrative Neuroimaging, University of Oxford, Oxford, United Kingdom; Research Department of Epilepsy, UCL Queen Square Institute of Neurology, University College London, United Kingdom; Centre de Recerca Matemàtica (CRM), Bellaterra, Spain; Center of Functionally Integrative Neuroscience, Department of Clinical Medicine, Aarhus University, Aarhus, Denmark

**Author notes:** Joint first authorship (Ryshum Ali, Olivier Burta, Chiara Rossi).

**Keywords:** MEG, dipole sign ambiguity, source reconstruction, beamforming, phase coupling

## Abstract

Source reconstruction is widely used to map magnetoencephalography (MEG) and electroencephalography (EEG) signals into localized brain activity, but the polarity of each reconstructed time course is arbitrary and differs across sources and subjects. This sign ambiguity compromises group-level analyses and any computed metric sensitive to the polarity of the time courses. Existing approaches to correct for this problem mostly rely on the event-related field (ERF), the trial-averaged response to repeated stimuli. As a result, no established alternative exists for sign-flipping in recordings without time-locked events, such as resting-state. Here, we propose SALSA (Sign Alignment for Source Analysis), a sign-flipping algorithm that can be applied to continuous MEG without task dependency. SALSA aligns polarities by maximizing the consistency of inter-regional phase coupling across subjects. Using controlled simulations with known ground truth, we show that SALSA recovers channel polarities introduced into synthetic MEG data. On task MEG data recorded during an auditory oddball paradigm, we benchmarked SALSA against existing sign-flipping approaches using group-level ERFs and a functional connectivity metric. We found that SALSA was the only approach to increase inter-subject phase consistency relative to data uncorrected for sign ambiguity. SALSA offers a systematically evaluated, task-independent approach to conduct sign alignment ahead of polarity-sensitive group-level analyses.

## Introduction

Magnetoencephalography (MEG) and electroencephalography (EEG) are non-invasive neuroimaging techniques that measure the electromagnetic activity resulting from the postsynaptic firing of cortical neuronal populations (Lopes da Silva, 2013). With a high temporal (milliseconds) and relatively high spatial resolution, and thanks to advances in source reconstruction methods, MEG has been brought to the forefront of research in neuroscience (Hämäläinen et al., 1993; Lopes da Silva, 2013; Westner et al., 2022). For instance, beamformers, one of the most widely implemented techniques, model the neural sources as dipoles and reconstruct the dipolar time courses with a spatial filter that weighs the contribution of each source to compose the sensors’ recordings (Westner et al., 2022).

However, when reconstructing source activity from sensor-level MEG data, the polarity of the resulting time courses is ambiguous. This poses an obstacle for group analyses, where each subject’s MEG data are source-reconstructed individually, and for recordings with multiple sessions per subject, as the same cortical source can be recovered with opposite signs. Averaging or concatenating across subjects can then attenuate or even cancel the shared neural activity, while also compromising connectivity and network analyses based on phase coupling between the time courses (Vidaurre et al., 2018).

The first source of sign ambiguity concerns the anatomical configuration of the neuronal sources in MEG. The postsynaptic activity of pyramidal neurons located in the cortical sulci forms the main contribution to the recorded MEG sensor data (Figure 1A), as these tangential currents generate magnetic fields that can be detected non-invasively outside the head (Hämäläinen et al., 1993; Lopes da Silva, 2013). The issue is that two sources with the same location and magnitude, one with positive polarity and a given orientation, the other with negative polarity and the opposite orientation, generate identical magnetic fields (see Figure 1B). Therefore, the polarity of the reconstructed time course cannot be uniquely determined from MEG measurements alone. Another source of sign ambiguity arises when projecting the three-dimensional reconstructed current dipole onto a single dimension representing the general activity of a voxel. This dimensionality reduction is conducted via eigenvalue decomposition, as in principal component analysis (PCA). The resulting components have an indeterminate sign, yielding two equivalent solutions of opposite polarity (see Figure 1C). Sign ambiguity may also arise in subsequent analysis steps: e.g., when applying a parcellation and using PCA to extract the component that captures the most variance among the time series of all sources within a given parcel. Altogether, during the source reconstruction process, the sign of the representative eigenvector is arbitrarily chosen for each subject and each session, leading to sign inconsistencies.

**Figure 1.**
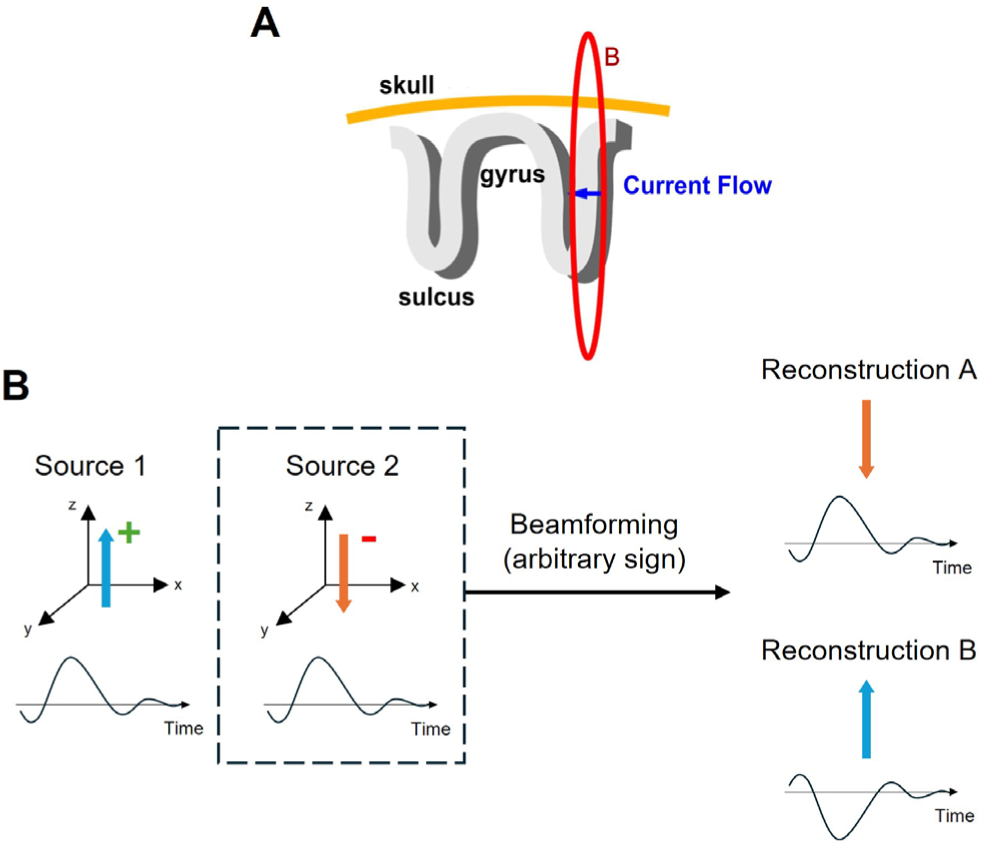
Sign ambiguity in source-reconstructed MEG. **(A)** Cortical gyri and sulci. The dark blue arrow represents the current dipole generated by neural activity, and the red circle denotes the associated magnetic field. MEG sensors mainly detect the magnetic fields generated by postsynaptic currents of pyramidal neurons in the sulci, whose orientation is tangential to the scalp. Illustration adapted from (Tamilia et al., 2017)**. (B)** Left: two dipoles (Sources 1 and 2), shown in the same coordinate frame, have identical magnitude and location but opposite orientation (colored arrow) and polarity (+ or − sign). Because the two reversals cancel, the MEG sensors pick up the same signal from both sources. Right: beamforming is applied to the measured MEG signal to estimate the activity of Source 2 (dashed box). Through eigenvalue decomposition, the three-dimensional dipole is projected onto a single axis. Either direction of this axis provides an equally valid solution (Reconstruction A or B), and the arbitrarily chosen direction determines the sign of the source-reconstructed time series.

Several neuroimaging signal processing toolboxes have proposed different techniques to overcome this sign ambiguity. Most of these methods are developed for event-related paradigms and use the event-related field (ERF), a waveform extracted from the continuous data that captures the modulation of the brain response following stimulus onset (Gramfort et al., 2013; OHBA Analysis Group, 2020). These methods assume that (i) the evoked response is temporally well-aligned across trials and has consistent polarity, and (ii) the magnitude of the evoked response at the trial level is sufficiently large relative to background activity.

A refinement of these ERF-based approaches was recently proposed by Marshall and colleagues, where PCA is applied to the trial-averaged ERFs of each brain region per subject (Marshall et al., 2022). The data are then sign-flipped for subjects with a negative first principal component. (Marshall et al., 2022). However, these techniques are limited by the fact that they require a temporally constrained task. Therefore, they are not applicable to resting-state data or more complex tasks that do not produce clear time-locked responses and, thus, cannot be treated via event-related analysis.

In this work, we describe a method to overcome sign ambiguity without requiring task-related information. Earlier versions of the method have been used in research articles (van Es et al., 2025b; Van Schependom et al., 2019; Vidaurre et al., 2016, 2018), but it has never been formalized or evaluated. Our algorithm assumes that the sign of the phase relationships between regions is consistent across all subjects (i.e., either in-phase or anti-phase). This is reasonable insofar as subjects are expected to share the same neurophysiological mechanisms. We present a Python package implementing the proposed algorithm and evaluate its performance on simulated data. Since most existing sign-flipping approaches rely on event-related responses, we also benchmark our algorithm against them using task-based MEG data.

## Methods

### 2.1. The SALSA algorithm

We aim to address sign ambiguity in source-reconstructed, parcellated, and leakage-corrected (see below) MEG by aligning source time courses to maximize inter-subject consistency in phase coupling across brain regions. Specifically, we ask for this relationship to be either consistently in-phase or anti-phase across subjects. The assumption here is that human brains should share a basic configuration of neural communication, due to shared anatomical and physiological features.

Based on this assumption, we propose SALSA (Sign Alignment for Source Analysis), a search algorithm that finds the optimal polarity of each source time course per subject by maximizing a defined objective function across multiple subjects. SALSA has three key components: (i) the summary statistic that represents the data: the autocorrelation (AC) tensor, which captures the lagged cross-correlations between all source pairs for each subject; (ii) the set of parameters representing the solution: the flips matrix, telling which channels are flipped for each subject; and (iii) the score metric that we aim to optimise: a scalar measure of inter-subject consistency in phase coupling given the AC and the flips matrix. We now describe how these three components are computed and how they come together to form the algorithm.

#### 2.1.1. Autocorrelation Tensor

Before performing the sign-alignment of source time courses, we construct an autocorrelation (AC) tensor for each subject separately. First, the input source-reconstructed data are standardized to zero mean and unit variance, computed separately for each of the *p* channels. For each subject, the data are then time-delay embedded. Given a set of lags *t* ∈ {−*m*, …, +*m*}, with *m* being the maximum lag parameter, this process yields a new “wider” matrix with *p*(2*m* + 1) columns, containing (2*m* + 1) lagged versions of every channel (for example, the first (2*m* + 1) columns of this matrix contain lagged versions of the first channel data) (Vidaurre et al., 2018). We then compute the cross-correlation between all *p*(2*m* + 1) lagged channel time series, producing a correlation matrix *C* ∈ ℝ*^p^*^(2*m*+1)^ ^×^ *^p^*^(2*m*+1)^.

Figure 2A shows how *C* can be decomposed into *p*^2^ sub-matrices *C_c_*_1_*_c_*_2_, one for each channel pair (*c*_1_, *c*_2_). From each off-diagonal sub-matrix *C_c_*_1*c*2_, whose entries reflect correlations between lagged versions of channels *c*_1_ and *c*_2_, we extract the anti-diagonal elements. These correspond to a fixed relative lag between the considered channel pair, directly capturing their phase relationship, and are used to populate the AC ∈ ℝ*^p^*^×*p*×(2*m*+1)^tensor (see Figure 2B). This procedure is repeated for all *n* subjects, yielding a 4D tensor (Figure 2C) that will be reused throughout the entire optimisation scheme of SALSA, with no further use of the original time series.

**Figure 2.**
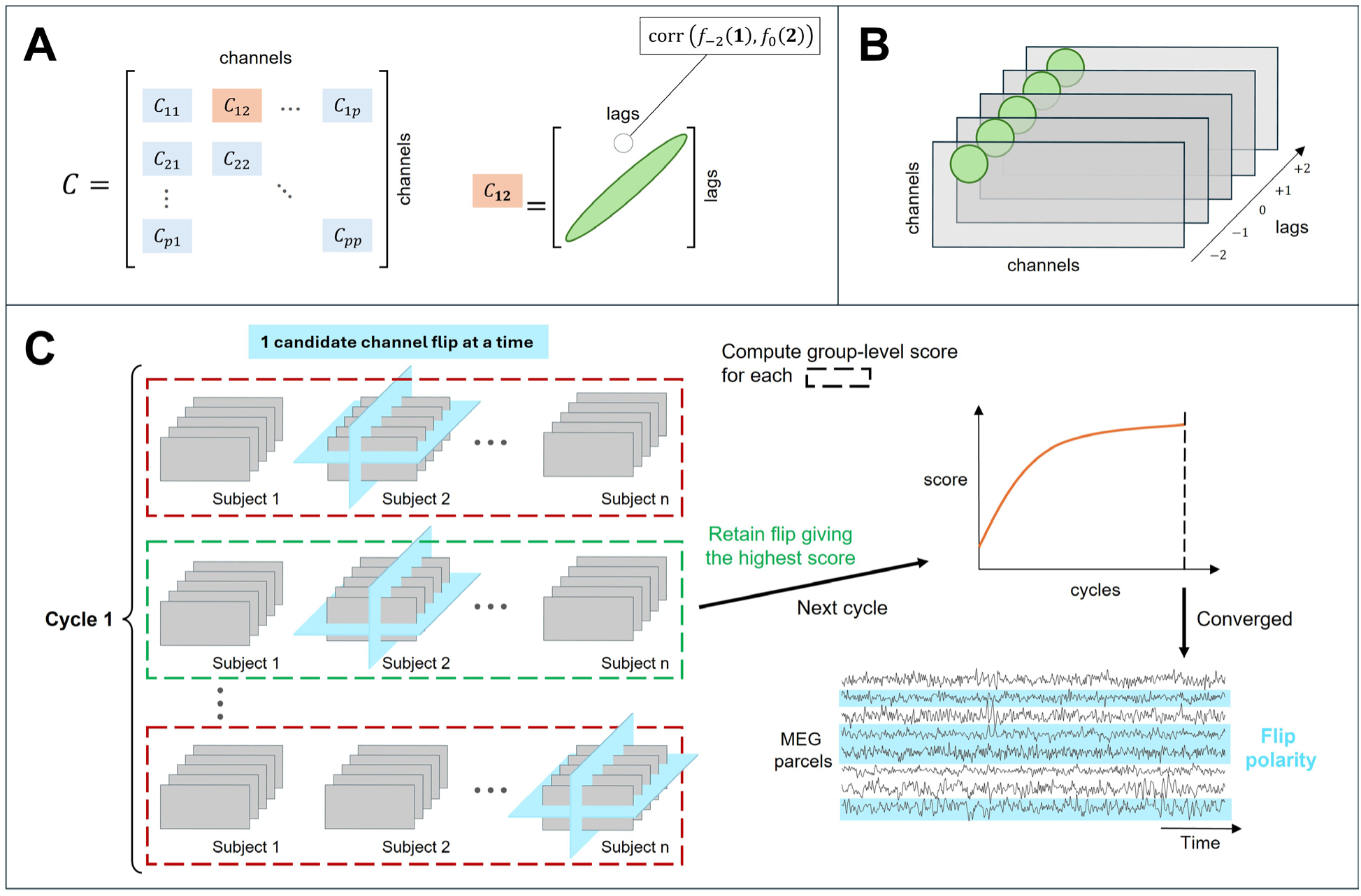
Flowchart of the SALSA algorithm. **(A)** Correlation matrix. For each subject, we compute the cross-correlation between all lagged channel time series. In this example, each channel is lagged using 5 lags, i.e. spanning {−2, …, +2}. After decomposing the correlation matrix *C* into sub-matrices, we extract the anti-diagonal elements (green) of, e.g. sub-matrix *C*_12_, capturing the phase relationship between channels 1 and 2. **(B)** Autocorrelation (AC) tensor. The extracted anti-diagonal elements populate the subject-level AC ∈ ℝ*^p^*^×*p*×(2*m*+1)^ tensor, one slice per lag. The entries of this tensor are computed once and, except for the sign, kept fixed throughout the subsequent optimization phase. (**C)** Greedy search to find the optimal flips matrix *F* that maximizes the phase consistency across subjects. At each cycle, all possible sign flips are evaluated one at a time. Flipping the polarity of channel *c* for subject *s* is equivalent to multiplying all AC tensor entries crossing the corresponding blue planes by−1. The candidate flip producing the highest score (green dashed box) is retained and *F* is updated accordingly. The algorithm iterates until the score improvement falls below a convergence threshold *δ*. Finally, the optimal polarities stored in *F* are applied to the original time series of each subject.

#### 2.1.2. Flips Matrix

Flipping the dipole polarities is performed using a flips matrix *F* ∈ {−1, +1}*^n^*^×*p*^, a subjects-by-channel matrix showing whether a channel’s polarity is to be flipped (−1) or not (+1). As illustrated in Figure 2C, an entry *F_s_*_,*c*_ = −1 corresponds to a sign flip applied to channel *c* within the AC tensor of subject *s*. The optimal *F* would be the one that, when applied to the 4D AC tensor, maximizes inter-subject consistency in phase coupling.

#### 2.1.3. Score Metric

To quantify how consistent the phase coupling is across subjects for a given *F*, SALSA uses a score, computed as the Frobenius norm of the mean AC tensor averaged across all *n* subjects:

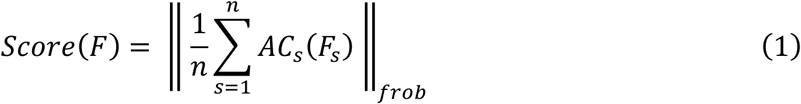

where ‖ ‖*_frob_* denotes the Frobenius norm and *AC_s_*(*F*) is the AC tensor of subject *s* after applying *F_s_*, the *s*-th row of *F*, to flip the polarities. This metric is large when the lagged cross-channel correlations, reflecting phase coupling, share the same sign across subjects.

#### 2.1.4. Optimization Strategy

To find the optimal *F* that maximizes the score in Equation 1, we employ a greedy search. At each iteration, referred to as a cycle, all possible subject-channel (*s*, *c*) combinations are evaluated by tentatively inverting, one at a time, the sign of *F_s_*_,*c*_ and computing the resulting score. The flip yielding the greatest score improvement is retained for the next cycle (see Figure 2C). If no flip leads to an improvement exceeding a threshold *δ*, the algorithm is considered to have converged. This procedure is repeated for multiple independent runs, each using a different random initialisation of *F*, where the channels to be flipped are drawn separately for each subject from a Bernoulli distribution with probability *ρ_init_*. For a total of *k* runs, the first run is always initialised with *F* as an all-ones matrix (i.e., no flips), while the remaining *k* − 1 runs each start from a different random initialisation of *F*.

All main results were obtained using the score metric described in Section 2.1.3, which guides the greedy search by evaluating inter-subject consistency across all channel pairs simultaneously (hence referred to as global). We also considered an alternative metric that assesses phase-coupling consistency separately for each channel pair (*c*_1_, *c*_2_). This pairwise implementation may be more sensitive to localized phase relationships between specific regions, but is computationally more expensive (equation in Supplementary Section A). Because the global and pairwise score metrics yield values on different numerical scales, applying the same convergence threshold *δ* may inadvertently favour one metric over the other by allowing more or fewer optimisation cycles within a run. To address this, we ran SALSA using both metrics across three threshold values *δ* ∈ {0.001, 0.0001, 0.00001}, the results of which are reported in Supplementary Section E.

#### 2.1.5. Hierarchical SALSA

For datasets with a larger number of subjects *n*, the computational cost of the greedy search grows substantially. Each cycle requires evaluating *n* × *p* candidate flips, and run convergence is expected to require more cycles since more subject-channel pairs need to be optimized. To reduce this heavy computational burden for large *n*, we implemented a hierarchical solution to SALSA. In particular, subjects are randomly divided into groups, where SALSA is then run independently on each group. The sign-aligned time series within each group are then concatenated across subjects to form a single “super-subject”, which is subsequently treated as an individual subject at the next level of the hierarchy. Unlike the standard SALSA implementation, where *k* runs are performed on all *n* subjects simultaneously, the hierarchical solution runs *k* independent initialisations within each group and at each level. For example, with a specified hierarchy of [10, 2] and *n* = 50, *k* runs are performed independently within each of the 10 groups at the first level, and *k* runs again within each of the 2 groups at the second level.

### 2.2. Existing Sign-Flipping Approaches

Three approaches proposed in the literature to resolve sign ambiguity are: the default approach implemented in MNE, a PCA-based approach, and a template-based approach implemented in OSL. The first two determine the optimal sign of each subject’s time series from the event-related field (ERF), while the last one can be applied to continuous recordings and uses a representative template subject to align the remaining subjects.

#### Default

We refer to the method implemented in the MNE software as the default approach, since it is the most straightforward way to resolve the sign ambiguity in event-related analyses (Gramfort et al., 2013). For each subject and region, the mean ERF waveform is computed across trials. The dominant polarity of each waveform is determined by the sign of the difference between its maximum and minimum peaks. For each region, we count how many subjects have the same dominant peak polarity, and we invert the polarity of the subjects forming the minority group.

#### PCA-based

Marshall et al. proposed a PCA-based approach to resolve sign ambiguity (Marshall et al., 2022). For each region, the mean ERF waveform is computed per subject across trials, and PCA is applied to the resulting subjects-by-time matrix. If a subject’s score on the first principal component is negative, that subject’s time course is sign-flipped. As with the default sign-flipping approach, this procedure is applied independently per region.

#### OSLE

The OSL Electrophysiological Data Analysis Toolbox (osl-ephys, OHBA Analysis Group, University of Oxford, UK) includes a functionality to correct for sign ambiguities (van Es et al., 2025a). The main idea is to select a template subject and to align the polarities of each remaining subject to it independently. For each subject, the parcellated MEG data are time-delay embedded (15 lags), standardized, and summarized by a single covariance matrix computed across all time-shifted channel copies. For all subject pairs, the Pearson correlation between the upper triangular of their absolute covariance matrices is computed. These similarities are summed for each subject, and the subject with the median summed similarity is selected as the template for sign-flipping. For each remaining subject, a stochastic hill-climbing search retains the flips that increase the Pearson correlation between its flipped covariance matrix and that of the template. The search is controlled by the number of independent initialisations, the number of iterations per initialisation, and the maximum number of channels flipped per iteration; we used the values recommended in the toolbox’s example scripts (*n_init_* = 3, *n_iter_* = 2500, maximum number of flips = 20). Throughout the results, this approach is referred to as OSLE.

### 2.3. Simulations

To evaluate our algorithm under controlled conditions, we applied it to synthetic source-reconstructed MEG data in which sign ambiguity was artificially introduced. The sign inconsistency is achieved by applying a ground-truth flips matrix *F*_0_ to the synthetic data, where each subject-channel pair independently has a probability of *ρ_amb_* to be flipped. Higher values of *ρ_amb_* introduce more polarity flips, hence increasing the difficulty of recovering the original sign configuration.

Synthetic data were generated using a multivariate autoregressive (MAR) model (Penny and Roberts, 2002). For a given number of channels *p*, we used a 15^th^-order MAR model pre-fitted to real source-reconstructed MEG data via maximum likelihood. Synthetic time series of 150,000 time points were then generated for each simulated subject by sampling from the fitted model with additive Gaussian noise. Separate model parameters were estimated for *p* ∈ {10, 25, 50} channels.

Since the ground-truth flips matrix *F*_0_ is known, we can directly assess how accurately the algorithm recovers the original sign configuration. In particular, we compare the ground-truth *F*_0_ against the optimal solution *F* provided by SALSA using a clique-based accuracy metric. For every unique pair of subjects (*s*_1_, *s*_2_) and channels (*c*_1_, *c*_2_), we form a clique ∈ {−1, +1}^2×2^ from *F* and *F*_0_, and compute the product of each clique’s four entries. A clique is then assigned an accuracy of 1 if the products obtained from *F* and *F*_0_ are equal, and 0 otherwise. The overall accuracy is the mean over all possible cliques, yielding a value between 0 and 1, where higher accuracy values indicate better recovery of *F*_0_ by SALSA.

### 2.4. Task MEG data

To evaluate how well SALSA resolves sign ambiguity in real MEG data, we compared it against existing sign-flipping approaches. Although SALSA can readily be applied to continuous recordings without task dependency, existing approaches to resolve sign ambiguity rely on the event-related field (ERF) and therefore require time-locked events. To enable a comparison with these existing methods, we used MEG data recorded during a task. We assessed each algorithm’s ability to resolve sign ambiguity at the group level using two measures that are sensitive to sign inconsistencies: the magnitude of the ERF, and the magnitude of a functional connectivity (FC) metric (the phase locking value). The latter builds on the same premise as SALSA: given the shared anatomical and physiological architecture, any pair of regions is expected to be either consistently phase-locked or anti-phase-locked across subjects. As this FC metric does not require time-locked events, it can also be applied to resting-state data.

#### 2.4.1. Dataset

The acquisition of the data used in this work was approved by the ethics committee of the University Hospital in Brussels (Commissie Medische Ethiek UZ Brussel, B.U.N. 143201423263, 2015/11). The dataset included 25 healthy participants, comprising 10 females and 15 males; all participants signed an informed consent form. The mean age per group (expressed in years, mean±SD) was 49.3±11.2 for the female group and 50.2±11.5 for the male group, with no significant difference between groups (Wilcoxon rank-sum test, *p* = 0.9). Additionally, the mean education level (in years, mean±SD) did not significantly differ between the female (14.9±1.9) and male (15.5±2.4) groups (Wilcoxon rank-sum test, *p* = 0.3). All participants underwent an MEG recording while performing a classical auditory oddball task, followed by an MRI acquisition within a median of 5 days (interquartile range: 2–10 days). During the task, participants listened to a series of tones comprising 80% frequent (1000 Hz), 10% distractor (500 Hz), and 10% target tones (1500 Hz). Participants were instructed to press a button with their right hand as soon as they heard the target tone. The task consisted of 400 tones, each lasting 1 s, followed by a 1.5 s inter-stimulus interval. MEG acquisitions were carried out at the CUB Hôpital Erasme (Brussels, Belgium) utilizing the 306-channel NeuromagTM TRIUX system (MEGIN Oy, Croton Healthcare, Helsinki, Finland). The MEG system was located inside a lightweight magnetically shielded room (MSR, MaxshieldTM, MEGIN Oy, Croton Healthcare, Helsinki, Finland). Electrocardiography (ECG) and electro-oculography (EOG) channels were recorded simultaneously with MEG, and the participant’s head shape fiducial points (nasion, left and right preauricular) were digitized using an electromagnetic tracker (Fastrak, Polhemus, Colchester, Vermont). The participants’ magnetic resonance images (MRIs) were acquired at the Universitair Ziekenhuis Brussel (Jette, Belgium), with a 3T Achieva scanner (Philips, Best, Netherlands). T1-weighted 3D MRIs were obtained with the following acquisition parameters (repetition time = 4.939 ms, flipping angle = 8°, field of view of 230 x 230 mm², 310 sagittal slices), yielding a 0.53 × 0.53 × 0.5 mm³ resolution. The MR images were coregistered with the MNI152 atlas.

#### 2.4.2. MEG processing

MEG data were acquired at a sampling rate of 1000 Hz and a band-pass filter between [0.1–330] Hz. After acquisition, we applied movement correction and MEGIN’s MaxFilter to perform signal space separation (temporal extension) and remove data components associated with sources outside the head (Taulu et al., 2004). The MEG preprocessing was conducted in the MATLAB 2020b environment, utilizing toolboxes from MATLAB and OSL, recalling functions from SPM12 (Wellcome Trust Centre for Neuroimaging, University College London) and Fieldtrip (Oostenveld et al., 2011).

##### Preprocessing steps

First, we applied RHINO to coregister the MEG sensor coordinates with the patient-specific MRI scan, utilizing the subject-specific fiducial points (OSL, 2018). This allowed the MEG acquisitions to be mapped to MNI152 space for accurate source reconstruction. After downsampling the data to 250 Hz, we then applied two filters: a notch filter at 50 Hz to remove powerline noise, and a band-pass filter covering [1–45] Hz. Next, artefact rejection was conducted in two steps. First, the AFRICA (artefact rejection via independent component analysis) algorithm implemented in OSL (OSL, 2017) was used to decompose the MEG recordings into 62 components, and those that correlated (*r* > 0.5) with ECG and/or EOG recordings were discarded. Second, a data segmentation process labelled noisy 1-second windows using an outlier standard deviation test across sensors. Before source reconstruction, MEG acquisitions were normalized per sensor type (gradiometer and magnetometer) via eigenvalue decomposition.

##### Source-reconstruction and parcellation

We performed source reconstruction via a linearly constrained minimum variance beamformer (LCMV) as implemented in OSL (Woolrich et al., 2011), using all sensor types (magnetometers and planar gradiometers). Each participant’s MEG session was individually source reconstructed using the full recording and considering a scalar beamformer with a single-shell forward model with a 5 mm dipole grid covering the cortical areas, coregistered in MNI space (Dale et al., 2000). This scalar beamformer projects each 3D dipole onto a single axis of maximum power via eigenvalue decomposition, a step that introduces sign ambiguity since the sign of the resulting eigenvector is arbitrary (Westner et al., 2022). Afterwards, the beamformed data were parcellated utilizing an atlas with 42 cortical regions (Rossi et al., 2023). The parcels’ time courses were extracted by taking the first principal component across the time series of the voxels contained within each region, adding another source of sign ambiguity. Finally, the parcels’ time courses were orthogonalized to discard zero-lag components via multivariate symmetric leakage correction (Colclough et al., 2015). This last step is important here, as the resulting time series are now uncorrelated at zero lag.

##### Sign-flipping using SALSA

SALSA is applied directly to the continuous parcellated MEG data, prior to epoching. We ran *k* = 10 runs with *m* = 10, *δ* = 0.001, *ρ_init_* = 0.25, on the MEG dataset containing *n* = 25 subjects and *p* = 42 parcels. The run achieving the highest gain value at convergence was used to sign-align the continuous parcellated MEG data, from which ERFs were subsequently extracted (see below).

#### 2.4.3. Validation based on Event-Related Fields (ERFs)

To assess how sign-flipping affects the consistency of task-evoked brain responses across subjects, we performed an ERF analysis on the parcellated MEG data. Epochs of [−0.2; 0.8] s around stimulus onset were extracted and individually baseline-corrected using the pre-stimulus period [−0.2; 0] s. Rather than averaging the epochs across trials to improve the signal-to-noise ratio (Hari et al., 2018; Luck, 2014), ERFs were estimated using a two-level general linear model (GLM). We performed the event-related analysis via GLM separately for each sign-flipping technique, as illustrated in Figure 3. As a reference, we also consider the case in which no sign-flipping is applied to the parcellated data (raw). The first-level GLM extracted subject-specific ERFs for the different components of the first-level design matrix, all related to the task conditions. The first-level design matrix for the oddball task includes four regressors (constant, frequent, target, and distractor), from which seven contrasts were computed: the constant, the three paradigm conditions, and the between-condition contrasts distractor versus frequent (DvsF), target versus frequent (TvsF), and target versus distractor (TvsD). The second-level GLM fitted the subjects’ first-level ERFs to infer the group-level ERF for each first-level contrast. At the group level, we then performed a maximum t-statistic permutation test (*N* = 10 000) to identify time points at which ERF amplitude significantly differed from baseline. We corrected for multiple comparisons via maximum statistics across regions, time, and GLM contrasts.

**Figure 3.**
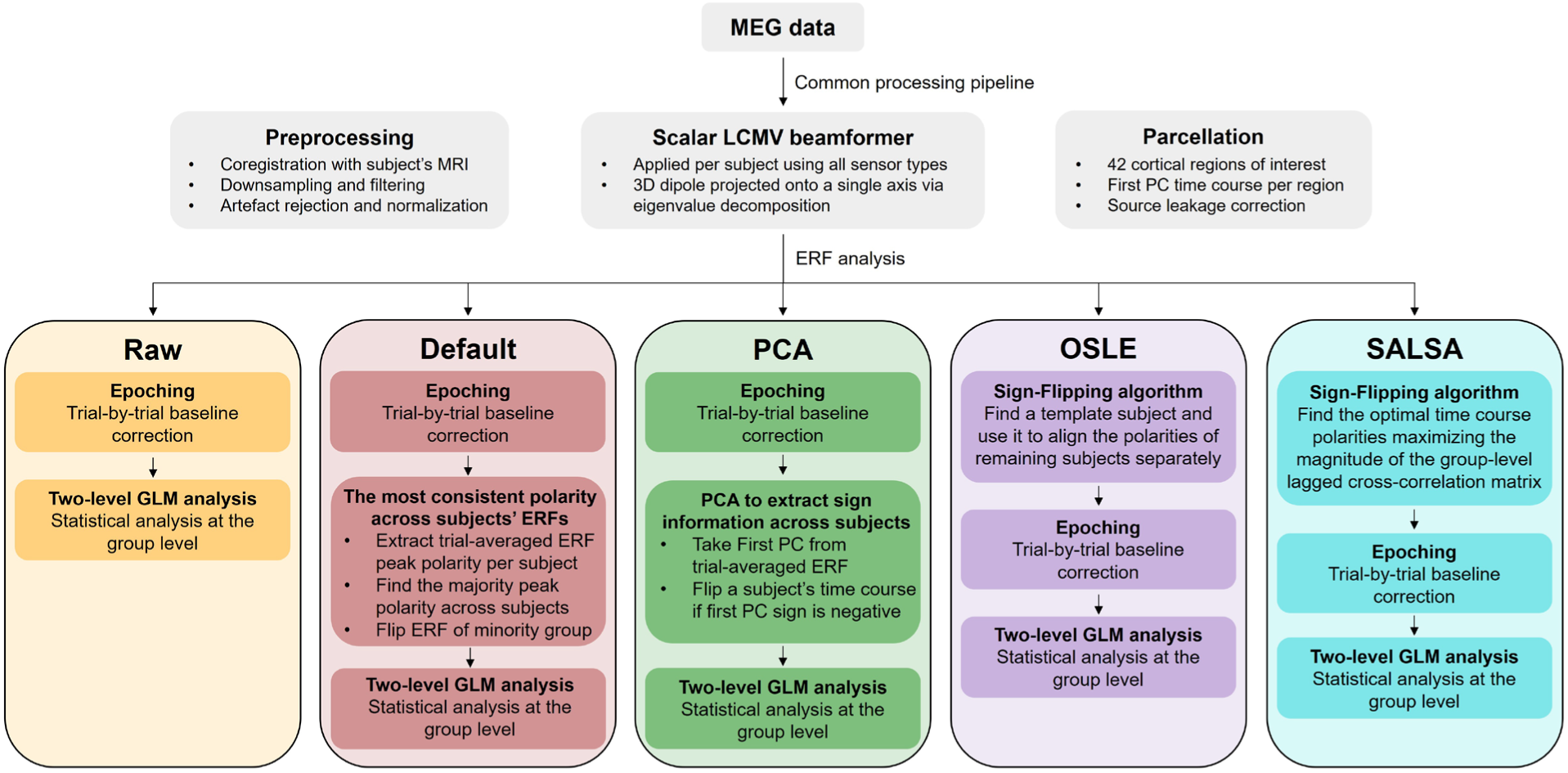
Different approaches to conduct sign-flipping on task-MEG data. All data pass through a common preprocessing pipeline including the source reconstruction. The following steps are then applied in different orders depending on the sign-flipping approach considered. Note that for the Default and PCA methods, the sign-flipping procedure is applied separately per region.

#### 2.4.4. Validation based on Functional Connectivity

To further assess sign-flipping quality in a way that can also be applied to resting-state, we looked at functional connectivity. In particular, we computed the phase locking value (PLV) as a measure of inter-subject consistency in phase coupling between brain region pairs. If two regions are functionally connected, their phase difference remains constant, yielding a higher PLV. Since PLV requires a single oscillatory component per region (Bruña et al., 2018), single-trial epochs were bandpass filtered in the alpha band [8–12] Hz before computing the PLV. We selected this frequency band as alpha-band activity has been associated with stimulus processing in auditory oddball paradigms (Barry et al., 2000; Öniz and Başar, 2009; Yordanova et al., 2001). Given that all subjects performed the same task, better sign alignment is expected to yield more consistent inter-subject phase relationships within this band. The PLV between channels *c*_1_ and *c*_2_ was computed as follows:

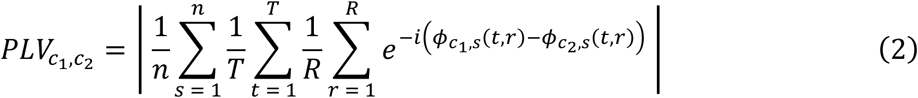

where *φ_c_*_1,*s*_(*t*, *k*) denotes the instantaneous phase of channel *c*_1_ for subject *s* at time point *t* in trial *r*, obtained as the argument of the analytic signal (computed via the Hilbert transform) of the bandpass-filtered epoch signal. Here, *n*, *T*, and *R* refer to the number of subjects, time points, and trials, respectively. Note that compared to the classical PLV, here the modulus is taken only after averaging across subjects, such that phase inconsistencies across subjects cancel out and produce a lower PLV.

The resulting PLV distribution (i.e., values pooled across all unique channel pairs) was compared across sign-flipping approaches (Default, PCA, OSLE, and SALSA) using a one-way ANOVA with Bonferroni-corrected post-hoc comparisons. The magnitude of PLV differences between sign-flipping approaches was quantified through Cohen’s *d*, comparing SALSA against each alternative approach (Default, PCA, and OSLE), including the parcellated data without sign correction.

## Results

We evaluated the performance and computational properties of SALSA in two complementary settings. First, we conducted simulations to assess the algorithm’s ability to recover ground-truth polarity flips applied to synthetic data under varying levels of problem complexity. Second, SALSA and other existing sign-flipping approaches were deployed on real task-MEG data. Since no ground truth is available for the real data, we used two different measures as evaluation criteria. The first is the event-related field (ERF), the same quantity that most of the existing approaches use to guide their sign-flipping. Sign inconsistencies across subjects attenuate group-level ERFs, so better sign alignment should recover more time points of significant evoked response magnitude. The second is functional connectivity, quantified as the phase locking value (PLV) between pairs of regions. Assuming that any pair of brain regions is consistently in-phase or anti-phase across subjects, better sign alignment should yield a higher PLV.

### 3.1. Simulations

We ran the SALSA algorithm on synthetic data for a varying number of channels *p* ∈ {10, 25, 50}, subjects *n* ∈ {5, 10, 25, 50, 100}, and ambiguous flip probabilities *ρ_amb_* ∈ {0.25, 0.50, 0.75}. For each configuration (*p*, *n*, *ρ_amb_*), the algorithm was run *k* = 5 times with different random initialisations of *F*, as described in Section 2.1.4. We repeated this procedure across 3 independent random seeds, each generating a different ambiguous dataset, yielding 15 runs total per configuration. The same 3 seeds were kept constant across all configurations, ensuring that variability in both the ambiguous data generated using *ρ_amb_* and the random initialisations of *F* was matched across configurations. The convergence threshold for all simulations was set to *δ* = 0.001, the initialisation probability to *ρ_init_* = 0.50, and the maximum lag to *m* = 10.

Figure 4A shows the sign-recovery accuracy across these configurations. Overall, the accuracy decreases with increasing *n*. For *p* = 10, the best run (i.e., the one converging with the highest score across *k* = 5 runs and 3 random seeds) reaches an accuracy of 1 even with increasing *n*. For configurations with more channels (*p* = 25, 50), however, the accuracy drops for large *n*, especially for *ρ_amb_* = 0.50. Interestingly, the accuracy results for *ρ_amb_* = 0.25 and 0.75 are highly comparable. This follows directly from the clique-based accuracy metric (see Section 2.3), which is invariant to negating the flips matrix, as *F* and −*F* yield identical accuracies. The run-to-run variability is reported in the Supplementary Materials (Section C, Figure S2). Across individual runs, the accuracy varies least when the number of channels is small (*p* = 10), especially when paired with few subjects. For larger *p*, accuracy across runs can range from 0.7–1, even for small *n*. The gap between the best run and the mean accuracy across runs increases with both *n* and *p*. Across all simulations, accuracy never fell below 0.6. More detailed results for a single SALSA run are reported in Supplementary Section B.

**Figure 4.**
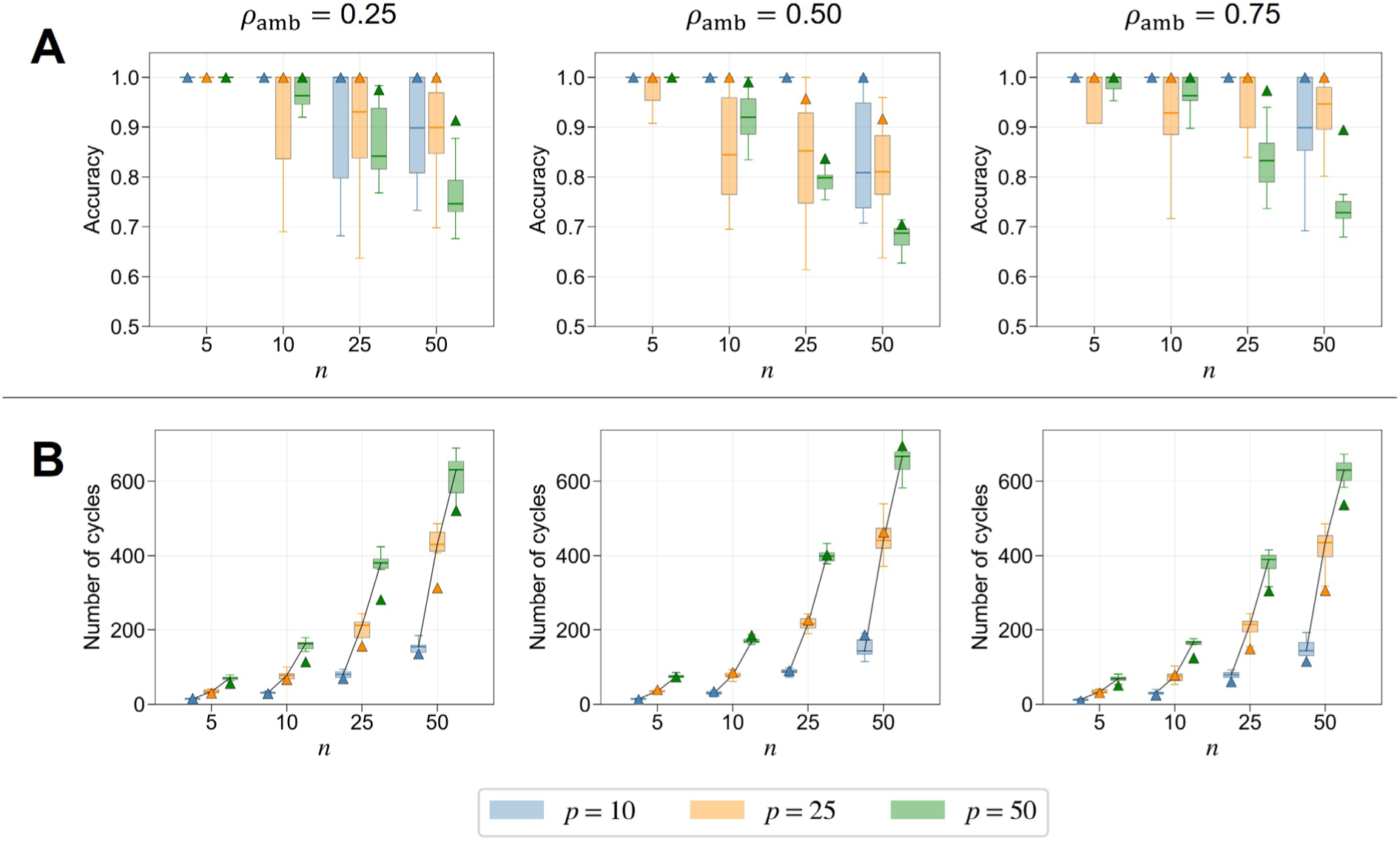
Performance as a function of dataset dimensionality and degree of sign ambiguity. The **(A)** accuracy of sign recovery and **(B)** number of cycles per run are shown for simulations with varying number of channels *p*, subjects *n*, and degree of ambiguity *ρamb*. Each column specifies the same *ρamb* across panels. For each configuration (*p*, *n*, *ρamb*), a boxplot summarizes the distribution of values across 15 runs (*k* = 5 initialisations and 3 random seeds), where the box spans the interquartile range (IQR), the central line indicates the median, and whiskers extend to 1.5 times the IQR. Triangles indicate the best-performing run per configuration. In panel B, the lines connect medians across parameters *p* within each *n*.

Figure 4B shows the number of cycles required for SALSA to converge. Overall, the cycle count increases with both *n* and *p*. For *ρ_amb_* = 0.25 (and similarly for *ρ_amb_* = 0.75), the best run consistently requires fewer cycles than the remaining runs within the same configuration. Interestingly, the same does not apply to configurations with *ρ_amb_* = 0.50. This prompted us to ask whether this behavior could be favored by a certain initialisation of *F*. We found that the first run (out of *k* = 5 runs), initialised with an all-ones matrix *F*, consistently produced the highest accuracy for configurations with *ρ_amb_* = 0.25. Specifically, this held in 100% of configurations for *ρ_amb_* = 0.25 but only 44.4% for *ρ_amb_* = 0.50, each proportion computed across the 36 configurations of (*p*, *n*, seed). When the first run did not produce the highest accuracy, the actual best run (i.e., one with a randomly initialised *F*) achieved a mean accuracy increase of 0.119 ± 0.095 over the first run. These results suggest that when sign ambiguity is higher, as expected in real MEG data, using multiple random initialisations of *F* becomes more beneficial. Besides the accuracy and number of cycles, we report the computational runtime of SALSA in Supplementary Section D.

### 3.2. Hierarchical SALSA

The standard SALSA implementation is expected to become computationally prohibitive when applied to datasets with large *n*. We therefore evaluated a hierarchical solution (see Section 2.1.5) on synthetic dataset configurations with *n* ∈ {50, 100} subjects, *p* ∈ {25, 50} channels, *ρ_amb_* = 0.50. Here, subjects were divided into 10 groups at the first level, and later into 2 groups of five super-subjects each. This procedure was repeated across the same 3 random seeds used in Section 3.1, where each seed simultaneously controls the ambiguous data generation and the random attribution of subjects into groups. This ensured that the hierarchical and standard SALSA solutions were evaluated on identical ambiguous datasets, allowing for a direct comparison of accuracy and computational runtime of both implementations. Since the hierarchical procedure returns one solution per synthetic dataset (rather than one per run), variability in performance was assessed across random seeds.

Figure 5 reports the accuracy and runtime of the hierarchical solution compared to the standard SALSA implementation, both evaluated on the same synthetic datasets with identical artificially introduced sign ambiguity. In terms of accuracy (Figure 5A), the hierarchical solution outperforms the standard SALSA implementation as *n* and *p* increase. Only for the configuration (*n* = 50, *p* = 25) does the standard implementation reach higher median accuracy across runs, albeit with greater run-to-run variability. The accuracy of the hierarchical solution remains consistently around 0.75 across all configurations, while the standard implementation degrades substantially with increasing *n* and *p*. The difference between the median accuracy reached by the two SALSA implementations is largest at (*n* = 100, *p* = 50).

**Figure 5.**
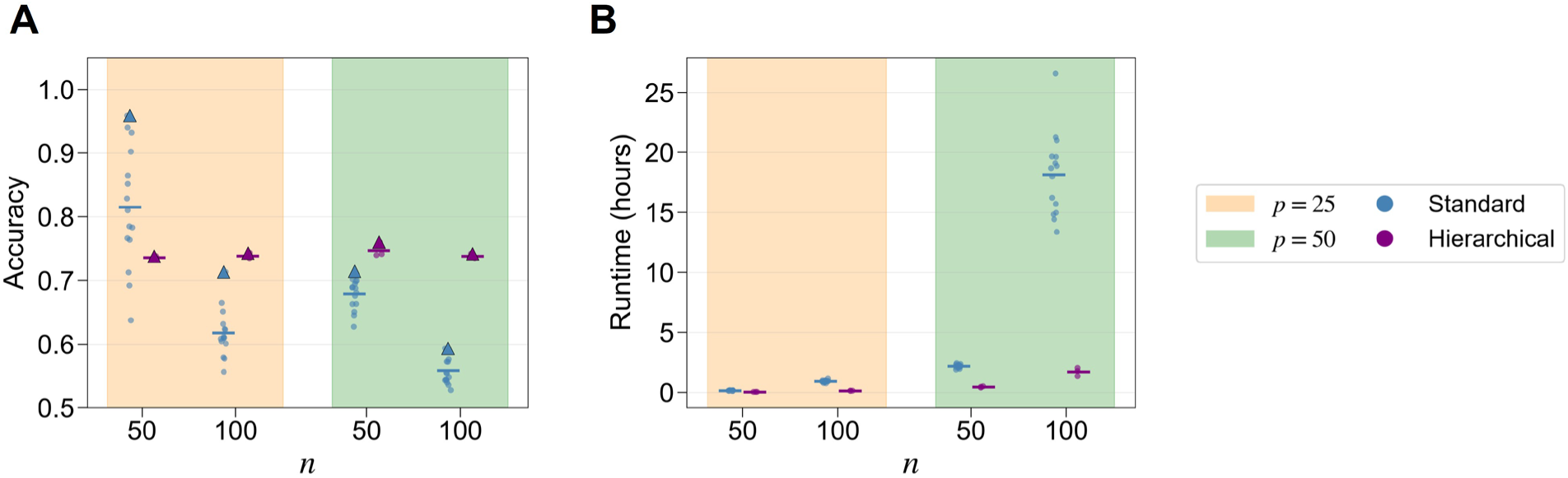
Hierarchical SALSA implementation for large datasets. We compare the **(A)** accuracy and **(B)** runtime expressed in hours using the standard (blue) versus hierarchical (purple) implementation of SALSA for simulated datasets with varying numbers of channels *p* and subjects *n*. Shaded areas specify a certain *p*. For each configuration (*p*, *n*), the horizontal line stands for the mean metric value across runs for the standard (15 runs; 5 initialisations and 3 random seeds) and hierarchical (3 random seeds) solutions, respectively. Triangles (panel A) indicate the best-performing run per configuration. All simulations were run on an Intel Core i9-12900F (12th Gen) with 128 GB of RAM, using a single core per run. Runtime was measured as wall-clock time.

Regarding the runtime to complete a single SALSA run (Figure 5B), the hierarchical solution is substantially faster than the standard implementation. The runtime difference becomes more important for *p* = 50, where the hierarchical solution is about 5.04 and 10.59 times faster for *n* = 50 and *n* = 100, respectively. Importantly, this speedup comes alongside higher accuracy in both cases. Had we picked a smaller convergence threshold *δ* for the standard SALSA runs, the algorithm could potentially go through more cycles before converging, likely reaching a higher accuracy. Because this would have led to unreasonable compute times on simulated datasets with large dimensionality, we instead assessed the effect of *δ* on the task-MEG dataset described in Section 2.4 (results reported in Supplementary Section E).

### 3.3. Performance on task MEG

#### 3.3.1. ERF magnitude

We first assessed how applying SALSA or alternative sign-flipping approaches affects group-level ERFs extracted from MEG data recorded during an auditory oddball task. Figure 6 presents the constant regressor (i.e., average activity across all paradigm conditions) and the contrasts of parameter estimates (COPES) for two representative cortical brain regions. To assess which sign-flipping approach best reveals differences between task conditions and their contrasts (e.g., target versus distractor stimuli), we also report the number of significant time points within the epoch window (see Figure 7).

**Figure 6.**
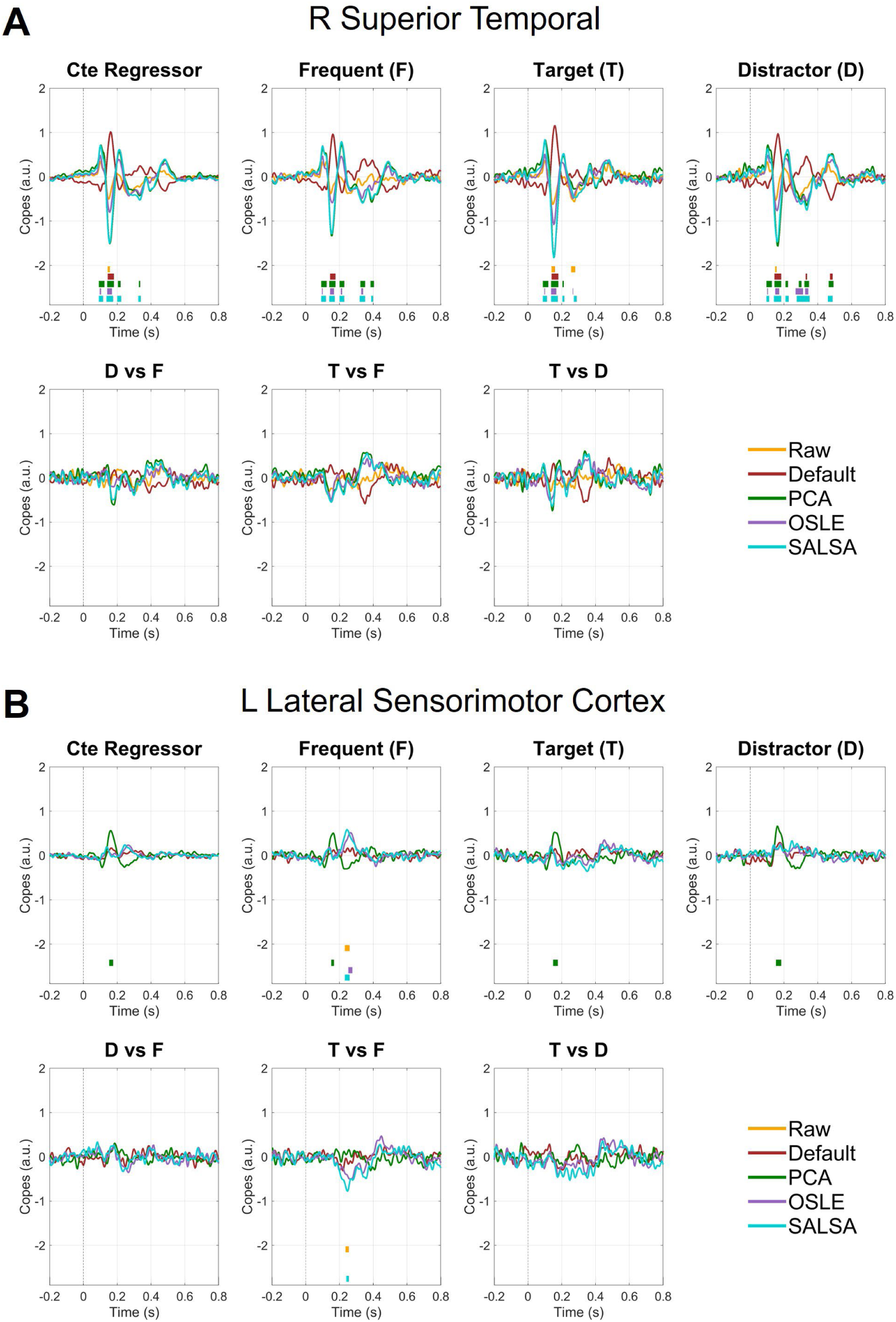
ERFs for different sign-flipping approaches. ERFs are shown for the **(A)** right superior temporal and **(B)** left lateral sensorimotor cortex regions. Each panel presents seven figures related to the ERF constant regressor (i.e., grand average), frequent (F), target (T), distractor (D), and their contrasts. In each subplot, we report the ERF resulting from the raw parcellated data (i.e., no sign-flipping) and four methods to overcome the sign ambiguity: default, PCA-based, OSLE, and SALSA. Horizontal lines under the curves highlight time points where the ERF waveform of the same color is significantly different from zero, hence the baseline level (max t-stat permutation, *N* permutations = 10 000, *p* < 0.025). The maximum statistic ensures adequate correction for multiple comparisons across time points, regions, and number of contrasts in the first-level design matrix. All y-axis values are in units of 10^−3^.

**Figure 7.**
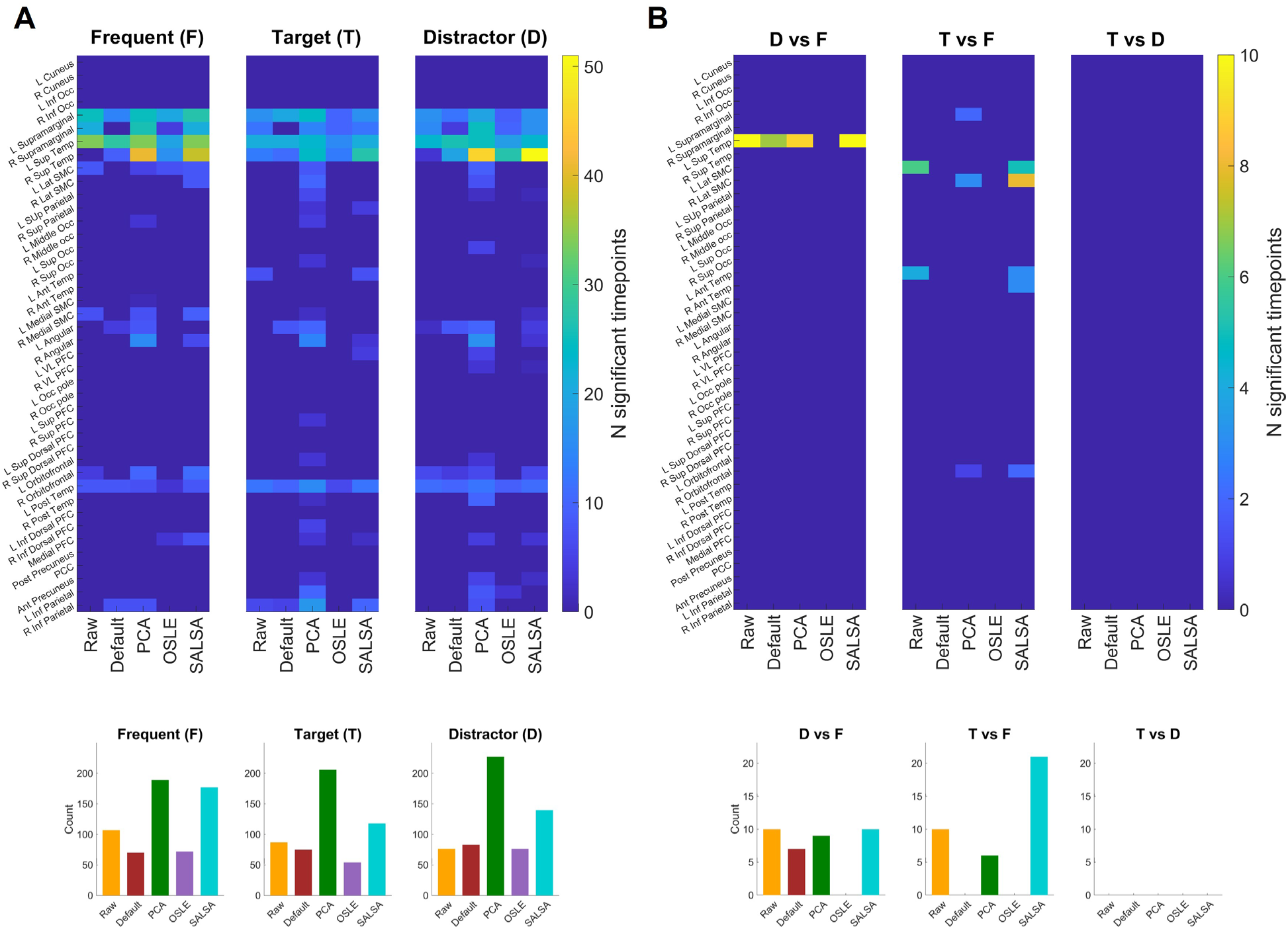
Number of significant ERF time points. Results are shown separately for **(A)** individual stimulus types (Frequent, Target, Distractor) and **(B)** contrasts between those task conditions. Within each panel, the top visual shows the number of time points (within the epoch window of [−0.2; 0.8] s around stimulus onset) where the ERF significantly differed from zero (max t-stat permutation, *N* permutations = 10 000, *p* < 0.025), separately per region (rows), sign-flipping approach (columns), and contrast (subplot). Color scales are kept separate for both panels such that differences between sign-flipping approaches can be visualized separately within the individual stimulus types (A) and within their contrasts (B). The bottom visuals within each panel show bar plots where the number of significant time points is summed across regions.

The default approach performs similarly to the uncorrected raw data, with stronger ERF peaks recovered in some regions (e.g., right superior temporal region, Figure 6A). The OSLE approach performs similarly to the default approach for individual stimulus conditions (Figure 7A), without improving on the results obtained from the uncorrected raw data. The PCA-based approach tends to better preserve the ERF’s oscillatory structure compared to the default and OSLE approaches. For individual stimulus conditions (frequent, target, distractor), PCA is often the only approach capable of revealing significantly modulated ERF waveforms (max t-stat permutation, *N* permutations = 10 000, *p* < 0.025) in a given region (Figure 7A). As shown in Figure 6B, the left lateral sensorimotor cortex (SMC) shows target and distractor peaks that only the PCA-based approach recovers, with none of the default, OSLE, or SALSA approaches detecting significant time points.

However, SALSA also recovers significant ERF modulations relative to baseline activity across multiple brain regions and task conditions. Notably, in the right lateral SMC (Figure 6B), SALSA uncovers a significant ERF modulation for the target versus frequent contrast that other approaches fail to capture. Overall, whereas the PCA-based approach most frequently identifies significant time points for individual stimulus conditions (Figure 7A), SALSA appears more effective at revealing significant time points for contrasts between task conditions (Figure 7B), mainly for target versus frequent. More specifically, summed across regions, PCA identified 622 significant time points across the three individual task conditions, compared with 435 for SALSA (Figure 7A), whereas across the three contrasts SALSA identified 31 compared with 15 for PCA (Figure 7B). Among the sign-flipping approaches applicable to continuous MEG recordings, SALSA was the only one to retrieve significant time points for contrasts between task conditions (Figure 7B), whereas OSLE retrieved none.

#### 3.3.2. PLV distributions

The second criterion we use for comparison is based on functional connectivity. It applies more broadly than the ERF assessment because it does not require a task and therefore does not necessitate a well-defined temporal alignment across trials. Specifically, we compared group-level PLV distributions across region pairs obtained after applying different sign-flipping approaches.

The results can be observed in Figure 8. A one-way ANOVA revealed a significant difference in mean PLV across the four sign-flipping approaches (*F*(3,3440) = 35.68, *p* < 0.001). SALSA achieved the highest mean PLV (mean ± std; 0.0120 ± 0.010), compared to the default (0.0085 ± 0.006) and PCA-based approach (0.0088 ± 0.006), both of which yielded lower mean PLV than the uncorrected raw data (0.0103 ± 0.008). The OSLE approach yielded a mean PLV of 0.0101 ± 0.009, similar to that of the uncorrected raw data (paired *t*-test, *p* = 0.60). Post-hoc comparisons confirmed that SALSA produced significantly higher PLV than the default (*p* < 0.001, Cohen’s *d* = 0.423), PCA-based sign-flipping (*p* < 0.001, Cohen’s *d* = 0.382), and OSLE approach (*p* < 0.001, Cohen’s *d* = 0.202). Notably, SALSA was the only approach to improve inter-subject phase coupling consistency compared to the raw data (Cohen’s *d* = 0.189), whereas the other approaches reduced it. We also verified that neither the choice of convergence threshold *δ* nor evaluating phase consistency separately per channel pair rather than across all channels simultaneously significantly affected downstream inter-subject phase coupling consistency (see Supplementary Section E).

**Figure 8.**
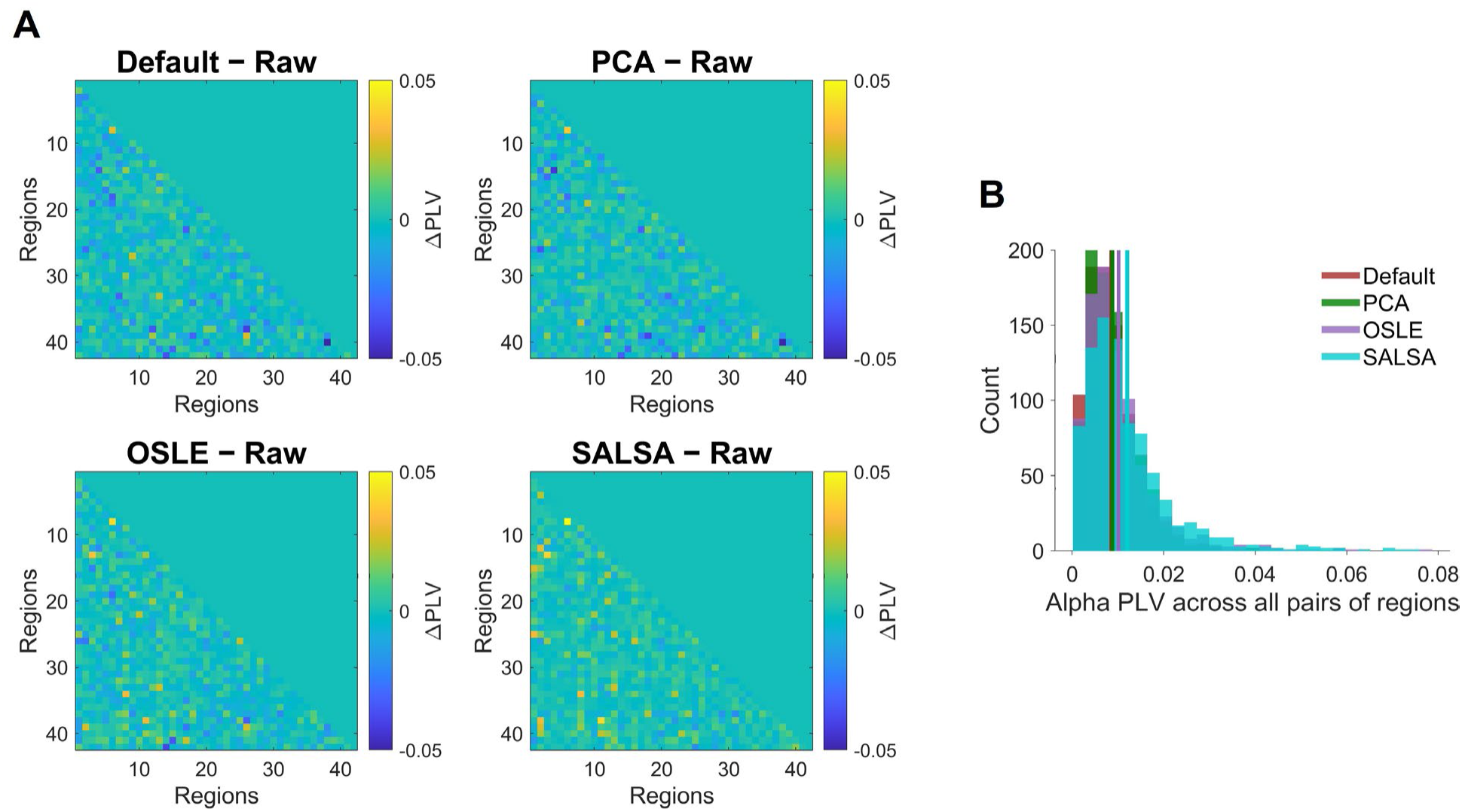
PLV distributions after applying different sign-flipping approaches. The PLV at each region pair was computed following Equation 2. **(A)** Functional connectivity (FC) matrices show ΔPLV, i.e., the difference between the PLV after sign-flipping and the PLV computed from the raw ERFs. Positive values of ΔPLV indicate improved inter-subject phase coupling for a given region pair relative to the non-sign-flipped raw data. **(B)** PLV distributions across all unique region pairs for each sign-flipping approach, with vertical lines indicating the mean across region pairs.

## Discussion

Reconstructing source activity from M/EEG is essential for studying brain function at the level of cortical regions. However, scalar beamformers arbitrarily assign a positive or negative polarity to each source time series. As this is done separately per subject, sign inconsistencies can attenuate shared neural activity, especially relevant in group-level analyses. Commonly used methods to correct for this sign ambiguity directly rely on the event-related potential (ERP) or event-related field (ERF), therefore requiring time-locked events in the form of task stimuli. As a result, resting-state MEG data (or less time-constrained task paradigms) currently lack a formalized and systematically evaluated method to resolve sign ambiguity. In this work, we propose SALSA, a sign-flipping algorithm that can be applied to continuous source-reconstructed MEG data that does not have a strong task dependency. Our approach aligns polarities by maximizing the consistency of inter-regional phase coupling across subjects. We provide SALSA as an open-source toolbox to perform sign-flipping ahead of group-level analyses.

Sign alignment matters most for analyses that use the signed source time series and combine data across subjects, such as phase-based connectivity or models fitted directly to concatenated time courses of all subjects. In contrast, the widely used analyses of spectral power or amplitude envelopes are insensitive to signal polarity. A prominent example of a group-level analysis that strongly benefits from sign alignment across subjects is the time-delay embedded hidden Markov Model (TDE-HMM) (Vidaurre et al., 2018). This data-driven approach identifies large-scale brain network dynamics from patterns of recurrent brain activity. The resulting HMM states are characterized by covariance structure estimated at the group level, such that inconsistent signs across subjects may reduce the model stability and the neurophysiological plausibility of the inferred states. Despite its importance for such analyses and the availability of existing approaches, sign-flipping appears to be an often overlooked step in the group-level analysis of source-reconstructed MEG (Dimitriadis et al., 2018; Siebenhühner et al., 2024; Zhang et al., 2022, 2021; Zhu et al., 2023). By providing an algorithm that does not require task events and is applicable to larger datasets through its hierarchical implementation, we hope that SALSA will encourage the research community to include sign-flipping as part of source-space MEG processing pipelines.

Among existing sign-flipping approaches, the default procedure in MNE (Gramfort et al., 2013) and a PCA-based approach (Marshall et al., 2022) both rely on the event-related potential (ERP) in EEG or event-related field (ERF) in MEG elicited by repeated stimulus presentation during a task. They thereby exploit the timing of the stimulus-locked response, information that SALSA does not use. This likely explains why, for individual task conditions (frequent, target, distractor), the PCA-based approach often found time points of significant ERP/ERF magnitude in regions missed by the other approaches. For contrasts between task conditions (e.g., target versus frequent stimuli), however, SALSA recovered more significant time points than the remaining methods. This dependence on stimulus timing can also become a limitation: approaches based on the ERP/ERF cannot be applied to resting-state data, and their performance is likely to degrade when brain responses are poorly aligned with external manipulations.

One existing approach that does not depend on task information is implemented in the osl-ephys toolbox (van Es et al., 2025a). It shares part of SALSA’s rationale, namely that the brain functional connectivity structure is expected to be common across subjects. This rationale has been applied in previous publications (van Es et al., 2025b; Van Schependom et al., 2019; Vidaurre et al., 2016, 2018), although never formally evaluated. The algorithmic implementations of both approaches, however, differ in a key aspect. Most importantly, sign-flipping from osl-ephys aligns each subject independently to a single template subject, so the outcome depends on the template and subjects are never optimized jointly, whereas SALSA evaluates all subjects together. To our knowledge, this existing algorithm has not been formally evaluated. When applying both algorithms to the task-MEG data, the template-based approach did not increase inter-subject phase consistency relative to uncorrected parcellated MEG data, while also yielding fewer time points of significant ERF magnitude compared to SALSA. Furthermore, our sign-flipping approach was the only one to improve the functional connectivity, as measured with the PLV, relative to the data uncorrected for sign ambiguity.

Beyond cryogenic MEG, SALSA can in principle be applied to any modality in which source reconstruction using beamformers yields time series with arbitrary polarity. This includes high-density EEG, where increased sensor density enables more accurate beamforming (Makkinayeri et al., 2025; Michel and Brunet, 2019; Murzin et al., 2011), and optically pumped magnetometer (OPM) MEG, whose wearable sensors enable recordings in naturalistic settings (Qin et al., 2026). In resting-state and naturalistic paradigms, where no repeated time-locked events are available, ERF-based approaches cannot be applied to resolve sign ambiguity. A recent OPM-MEG study of naturalistic recordings used beamforming for source reconstruction in 22 participants recorded in two sessions each (Gibson et al., 2026). Its reported downstream analyses used spectral power and amplitude envelope correlations, so a sign alignment step was not required. Such a dataset nevertheless involves two levels at which sign ambiguity can arise, across subjects and across sessions of the same subjects. Should polarity-sensitive group-level analyses be applied to these data, such as phase-based connectivity or a TDE-HMM, sign alignment with SALSA after source reconstruction and parcellation would be desirable. Although naturalistic recordings can be much longer than task recordings, the AC tensor required for SALSA is computed only once per subject and stored thereafter, so the duration of the greedy search itself does not depend on MEG recording length.

SALSA aligns polarities by maximizing the consistency of phase coupling across subjects, which frees it from task events and lets it use all subjects jointly. This design entails two main limitations. First, it rests on the assumption that phase relationships between regions are consistent across subjects. Our functional connectivity validation with the PLV quantifies the property that SALSA optimizes, so the higher PLV obtained with SALSA is partly expected by construction, which is why we additionally compared ERFs. Whether the assumption holds in cross-sectional datasets that pool different populations, such as patients and healthy controls or different age groups, although expected, remains to be assessed. Second, the greedy search evaluates every candidate flip in each cycle, so its cost grows with the number of subjects and channels. The hierarchical implementation mitigates this: its accuracy was lower than the best standard runs in smaller simulated datasets (e.g., 50 subjects and 25 channels), but for 100 subjects and the same number of channels, it was both more accurate and about tenfold faster.

In summary, SALSA provides a task-independent framework for aligning the polarity of source-reconstructed MEG time courses across subjects by maximizing the consistency of inter-regional phase coupling. In controlled simulations, it recovered introduced polarity flips with high accuracy. On task MEG data, it was the only approach to increase inter-subject phase consistency relative to uncorrected data, while the ERF-based approaches remained advantageous for individual task conditions in the auditory oddball. Together with its hierarchical implementation scalable to larger datasets, this positions SALSA as a practical step for group-level analyses of source-space MEG that do not rely on time-locked events.

