## Supplementary Materials for "SALSA: resolving sign ambiguity in M/EEG source reconstruction with no task dependence"

#### A. Pairwise score metric

As an alternative to the global metric (see Equation 1), the pairwise score metric is computed as follows. For each subject $s$, the lag vector $v_{s_1,c_1c_2} \in \mathbb{R}^{(2m+1)}$ is extracted from the AC tensor for each off-diagonal channel pair $(c_1, c_2)$ and z-scored to give $\tilde{v}_{s_1,c_1c_2}$. The score is then the average over all dot products $\langle\ \rangle$ between the normalized lag vectors across all unique subject- and channel pairs:

$$Score(F) = \frac{1}{N} \sum_{c_1 < c_2}^{p} \sum_{s_1 < s_2}^{n} \langle F_{s_1,c_1} \cdot \tilde{v}_{s_1,c_1c_2}, F_{s_1,c_1} \cdot \tilde{v}_{s_2,c_1c_2} \rangle \tag{3}$$

where $N = \binom{n}{2} \cdot p(p-1)$ the total number of terms. Alongside the main analyses conducted using the global metric (see Equation 1), we ran SALSA on the real task-MEG data using this pairwise metric (see Supplementary Section E).

#### B. Example of a single SALSA run

Below in Figure S1, we show some results of a single standard SALSA run on synthetic data ($p = 50, n = 100, \rho_{amb} = 0.50$). Most notable is the last subplot, where we can see that SALSA tends to flip channel X in a single subject and, during consecutive cycles, flips the same channel X for a decent subset of the remaining subjects.

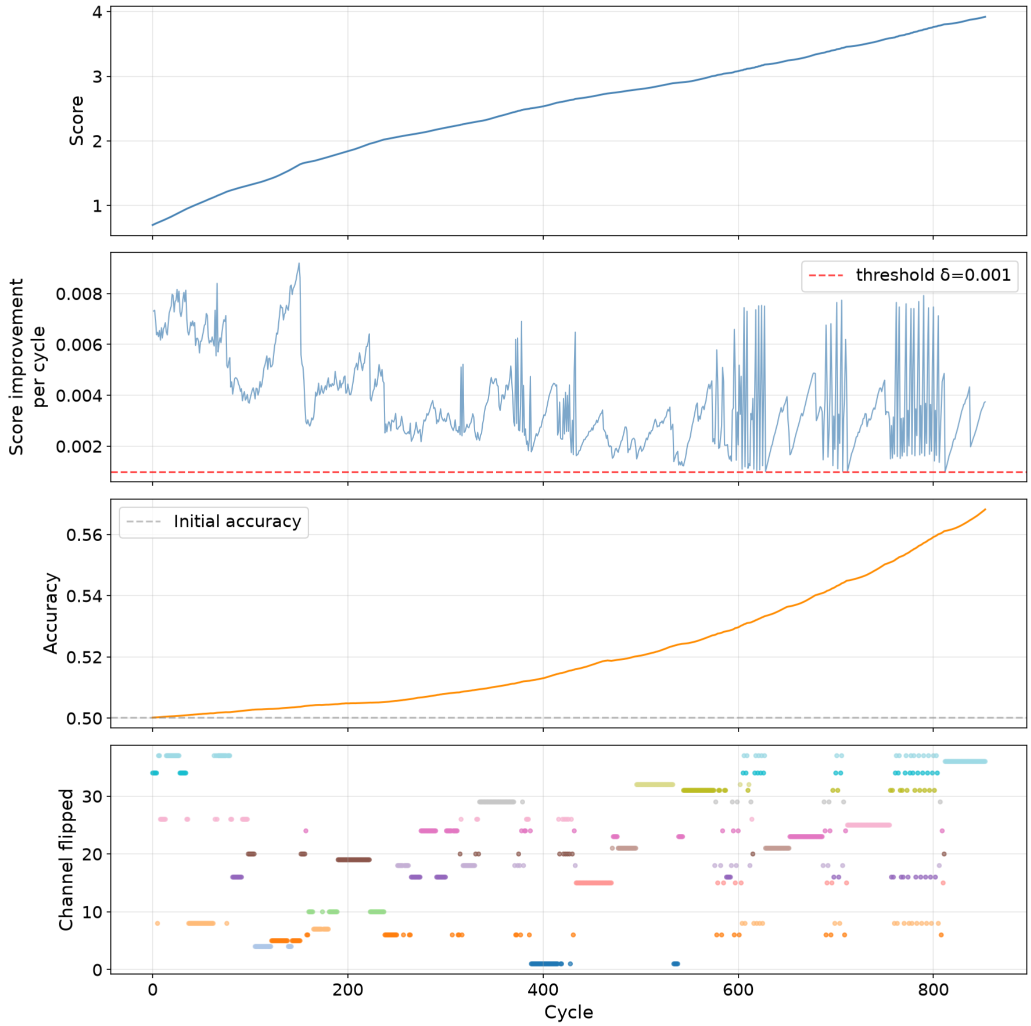


**Figure S1 Performance of a single SALSA run.** We ran SALSA on synthetic data ($p = 50, n = 100, \rho_{amb} = 0.50$), and show, in order from top to bottom: the score, the score improvement per cycle, the accuracy of group-level sign recovery, and which channel was flipped, always across cycles. The run converges when the score improvement per cycle falls below the convergence threshold $\delta$.

### C. Variability across runs

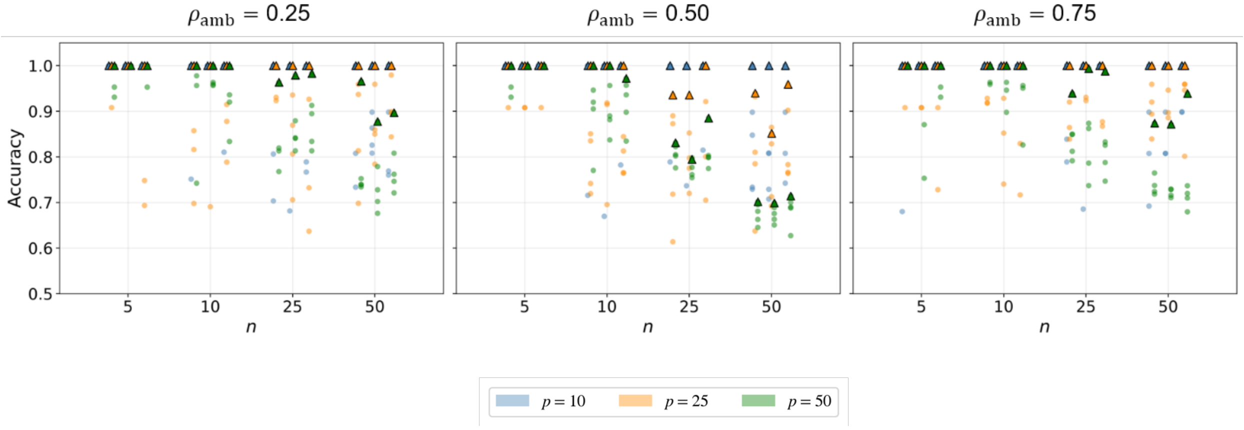


**Figure S2 Run-to-run variability in performance as a function of dataset dimensionality and degree of sign ambiguity.** The accuracy of sign recovery is shown for simulations with varying number of channels $p$, subjects $n$, and degree of ambiguity $\rho_{amb}$. For each configuration $(p, n, \rho_{amb})$, each tick on the x-axis is divided into 3 vertical lines, one for each random seed (which controls both the ambiguous data generation using $\rho_{amb}$ and the $k = 5$ initialisations of $F$). Triangles indicate the best-performing run per configuration and per random seed.

### D. Computational cost for standard SALSA

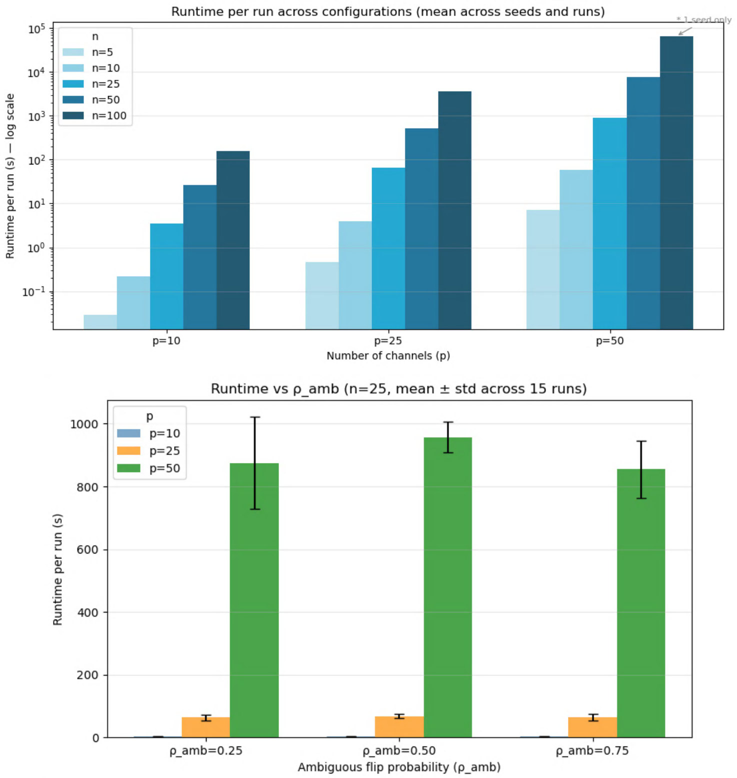


**Figure S3 Computational cost for standard SALSA runs.** All simulations were run on an Intel Core i9-12900F (12th Gen) with 128 GB of RAM, using a single core per run. Runtime was measured as wall-clock time.

### E. Effect of convergence threshold

We applied SALSA (for $k = 10$ runs, each with different initialisations of $F$) to the MEG data recorded during an auditory oddball task. From Figure S4, we observe that the first run out of $k = 10$ runs (i.e., the one initialised with an all-ones matrix $F$): starts with the highest score at cycle 0, converges to the highest score value, and does so in the least number of cycles, regardless of the score metric (global/pairwise) used. For the remaining runs (indexed 2–10), the effect of convergence threshold $\delta$ on the score progression differs depending on the score metric employed. For the global metric, lowering $\delta$ enables meaningful additional score improvement, but the first run remains far ahead regardless of how small $\delta$ is chosen. For the pairwise metric, however, lowering $\delta$ allows the runs with random initialisations of $F$ to catch up to the score value achieved by the first and best-performing run, albeit at the cost of substantially more cycles. Additionally, a run using the pairwise score metric takes about 15 times longer (in terms of compute time) to complete compared to using the global metric with the same $\delta$. This makes the pairwise metric computationally infeasible for larger datasets.

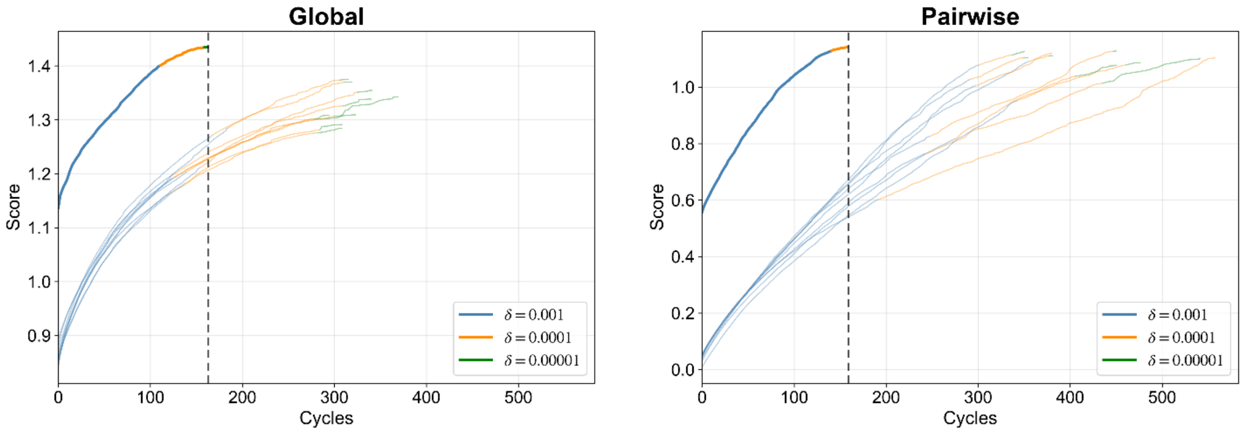


**Figure S4 Effect of lowering the convergence threshold $\delta$ on the score trajectory.** We ran the standard implementation of SALSA for the global (left) and pairwise (right) score metric for different values of $\delta$. Because the ambiguity generation and random flips initialisation were kept constant, each curve (corresponding to one of $k = 10$ runs) has three segments of three colors, The score trajectory for the largest threshold (blue) is identical to the first part of the trajectories for the smaller thresholds, so each later segment (orange, green) continues the previous one: lowering the threshold $\delta$only allows the search to run for additional cycles after the point where the larger threshold would have stopped. The bold curve marks the run with the highest final score, and the dashed vertical line marks the cycle at which it converged. The two panels use different y-axis ranges, as the global and pairwise scores are on different scales.

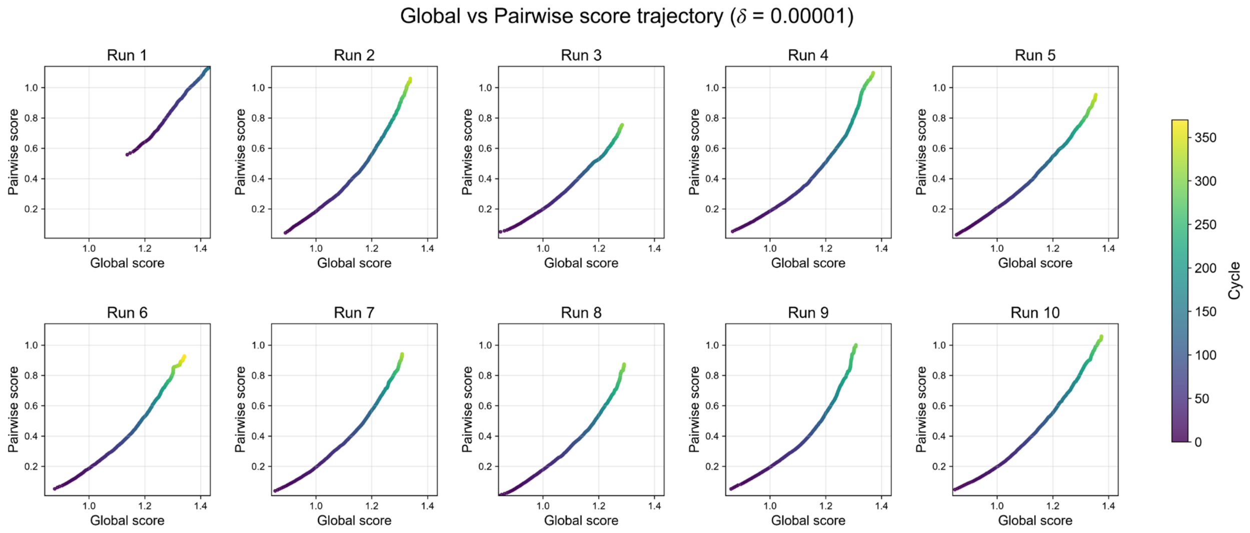


**Figure S5 Global versus pairwise score trajectory.** For each $k = 10$ run, we separately plot a scatter of the global score and pairwise score, where each dot stands for a cycle (color scale). For the same run, the sign ambiguity introduced and the random flips initialisation were kept constant. Across runs, only the ambiguous dataset generation changes. Run 1 is always initialised with an all-ones matrix (see Section 2.1.4).

To assess whether the choice of score metric and convergence threshold $\delta$ affected downstream PLV consistency across subjects, we ran SALSA on the task MEG dataset using both the global and pairwise score metrics across three threshold values. PLV distributions are shown in Figure S6. The global metric showed no significant difference in PLV distribution when the convergence threshold was lowered from $\delta = 0.001$ to $\delta = 0.00001$ ($t = -0.970, p = 0.332,$ Cohen's $d = -0.014$). The same held for the pairwise score metric ($t = -0.661, p = 0.509,$ Cohen's $d = -0.008$). Comparing the two metrics at the same threshold ($\delta = 0.001$) also revealed no significant difference ($t = 0.166, p = 0.868,$ Cohen's $d = -0.003$). Even when the pairwise metric was given a more permissive threshold ($\delta = 0.00001$) to compensate for its more constrained score range, no significant difference with the global metric at $\delta = 0.001$ was found ($t = 0.181, p = 0.858,$ Cohen's $d = 0.003$).

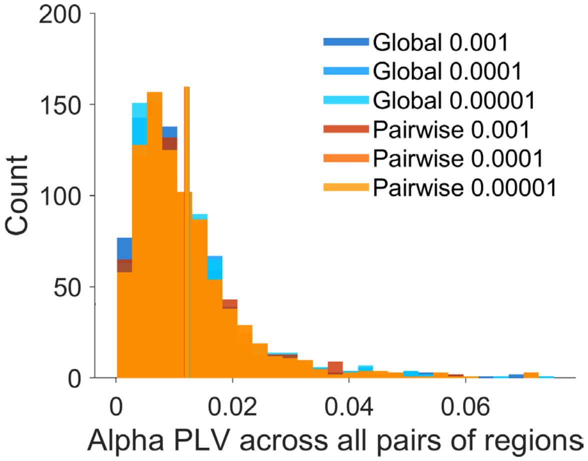


**Figure S6 PLV distributions obtained after applying SALSA for different score metrics and convergence threshold values.** PLV distributions across all unique region pairs for each metric-threshold combination, with vertical lines indicating the mean across region pairs. For each comparison (between 6 conditions, given 2 score metrics and 3 convergence thresholds), we ran paired *t*-tests across region pairs. Reported *p*-values are uncorrected, with the Bonferroni-corrected significance threshold for 15 comparisons being $p < 0.0033$.
